# Disruption of emotional processing by GBR 12909 in male and female mice highlights novel behavioral paradigms relevant to bipolar disorders

**DOI:** 10.1101/2025.10.02.679191

**Authors:** Maelle Certon, Bruno Brizard, Catherine Belzung, Alexandre Surget, Arnaud Tanti, Solal Bloch

**Affiliations:** Université de Tours, INSERM, Imaging Brain & Neuropsychiatry iBraiN U1253, 37032, Tours, France

## Abstract

Bipolar disorders (BD) are defined by a chronic recurrence of manic and depressive phases. Along with mood, acute phases are associated with altered emotions. The biological underpinnings of these changes are unresolved, mostly because modeling the cycling nature of BD is still a major challenge in preclinical studies. One model is based on GBR 12909 administration, a dopamine transporter inhibitor aiming at mimicking some dimensions of mania. It has recently been shown that this model generates a mixed phenotype with negative hedonic biases and anxiety along with hyperlocomotion. These studies have been only performed in male animals, and other behavioral dimensions relevant for BD remain to be explored, in particular recognition of conspecifics emotions and reactivity to danger. The objective of this study is to further characterize the GBR model in both sexes by introducing two novel behavioral assays, the sweeping/looming disk and the negative emotion recognition tasks to evaluate response to threat and emotion discrimination. First, we replicated the previous results in the GBR model: higher anxiety, hyperlocomotion, anhedonia in males. These phenotypes were less pronounced in females. GBR also induced a hypersensibility to threat in both sexes in the sweeping/looming disk. GBR abolished preference for the emotional target only in males, suggesting altered emotion recognition. This work introduces new phenotypic dimensions relevant to study BD and highlights the necessity to study both sexes which are not strictly equivalent in their behavioral responses.

## 1. Introduction

Bipolar disorders (BD) are severe neuropsychiatric conditions that affect over 40 million people globally. They are an important source of psychosocial impairment and reduce patient longevity by at least 10 years due to multiple comorbidities and the high risk of suicide (McIntyre et al., 2020). BD are characterized by the alternation of manic and depressive phases (or a combination called mixed states) interspaced by periods of remission (euthymia). Mania corresponds to abnormally increased mood, energy, motor activity, risk taking, irritability and decreased sleep, while depressive states show opposite symptoms in particular decreased mood and energy along with increased sleep (Vieta et al., 2018). Manic symptoms are a hallmark of the disease and need to be present to some extent (full mania, less pronounced hypomania, or mixed state with manic symptoms) to allow diagnosis, which partly explains the considerable delay to establish diagnosis (5 years on average). Multiple mechanisms have been involved in BD pathogenesis (Ashok et al., 2017; Balaraman et al., 2015; Chaves-Filho et al., 2024; Gonzalez, 2014; Kato, 2017; Sigitova et al., 2017). Nevertheless, there is no consensus towards a complete biological explanation of the origin and expression of the disease. BD are generally considered to have a similar expression in women and men (Kawa et al., 2005; Suominen et al., 2009). However, this question has received little attention compared to other pathologies such as depression and anxiety disorders (Bangasser and Cuarenta, 2021). Some studies hint at differences in comorbidity profiles and neuroanatomy, but more patients and studies are needed to explore the question of sex differences in BD (Jogia et al., 2012).

Due to the symptom heterogeneity and the complex cycling nature of the disease (Guglielmo et al., 2021), modeling BD in animals has been particularly challenging. Current models have been mainly focusing on manic-like phenotypes (Chaves-Filho et al., 2024) measured in behavioral tests traditionally used to identify depressive-like phenotypes, the expecting result in a mania model being the opposite (e.g., hyperhedonia, hypoanxiety, active stress coping strategies (Thomas et al., 2007, increased exploration and locomotion). In addition to these ‘classical’ mood-related phenotypes, emotional processing is one of the most altered behavioral dimensions in BD patients and is directly linked to manic and depressive symptoms (Bigot et al., 2020). Indeed, manic patients typically display positive emotional biases (Schönfelder et al., 2017) resulting in increased reward seeking and a decreased perception of danger, while negative biases generate the opposite outcome during depression (Bilderbeck et al., 2016). Patients also have disrupted emotion recognition and empathy responses both in euthymia and during acute phases (De Prisco et al., 2023; Derntl et al., 2012; Seidel et al., 2012). Some studies suggest a mood-incongruent deficit of facial emotion recognition (harder to recognize negative emotions in mania and positive emotion in depressive phase, Chen et al., 2006; Liu et al., 2024). Finally, episodic memory impairments have been described in BD (Cotrena et al., 2020; King et al., 2013) but have been little investigated in BD models and never in the GBR one. It is therefore crucial to implement novel behavioral tests reflecting these alterations of emotional responses including emotional biases, responses to threat, memory performance and emotion recognition to improve the validity and the characterization of BD animal models.

One way to model manic symptoms in animals is the injection of stimulants, typically amphetamine. This has some construct validity because dopaminergic system dysfunctions have been involved in BD (Ashok et al., 2017). Nevertheless, amphetamine affects other monoamine systems directly and triggers phenotypes not related to BD (e.g. stereotypies). GBR 12909 (GBR) is a more recent and valid stimulant model for BD. Compared to amphetamine, GBR injection increases risk taking, exploration and does not induce stereotypies (van Enkhuizen et al., 2015; Young et al., 2010). It is also highly specific for the dopamine transporter (Andersen, 1989). A recent study using the GBR mouse model has demonstrated increased negative biases in gustatory and olfactory preference tests along with increased locomotion, anxiety and combativeness (Bigot et al., 2022). Therefore, the GBR model may lead to both manic and depressive characteristics, which is closer to mixed states. These results highlight the importance of incorporating diverse behavioral dimensions to fully characterize an animal model of psychiatric disorders, and particularly of BD. To our knowledge, female animals have never been tested in this model, and are almost absent of current BD preclinical research.

In summary, assessing phenotypes beyond classical tests used in models of mood disorders is an important step towards further characterizing models of psychiatric disorders and in particular BD. We focused on the mixed manic-like phenotype induced rapidly by GBR injection. Hence, this work aimed at validating novel behavioral assays to improve model characterization in BD research. It also intended to complete the GBR model characterization building on the recent study on hedonic bias (Bigot et al., 2022), and investigate sex differences in this model. We performed five behavioral tests each corresponding to a behavioral dimension: anxiety (light dark box, LDB), coping strategy in response to a threat (sweeping/looming disk, SLD), hedonic bias (sucrose/quinine preference, SQP), long-term memory (novel object recognition, NOR), and emotion discrimination (negative emotion recognition in a conspecific, NER). The principles of SLD and NER have been published previously (De Franceschi et al., 2016; Scheggia et al., 2020) but have never been applied to BD research and rarely in neuropsychiatry in general. SLD consists in presenting threatening visual stimuli on a screen on top of the arena and recording the mouse reaction. The stimulus is either a small disk sweeping across the screen (mimicking a hovering predator) or a disk increasing in size until taking the whole screen (diving predator). This elicits two types of behavioral responses: flight (to a shelter) and/or freeze (De Franceschi et al., 2016). Typically, an immediate threat triggers flight (looming) while a distant one elicits freezing (sweeping). The SLD has the advantage of presenting an ecologically relevant situation to the mouse. It is also highly valuable for BD research because it reflects dimensions such as reactivity and vigilance to a stimulus (e.g. hypervigilance in mania), risk taking, and coping strategy choice. NER is a simple modified social preference task in which the experimental animal chooses to approach a previously stressed or a control mouse. In control conditions, the stressed mouse will be preferred by the experimental mouse (Scheggia et al., 2020). Given the empathy and emotion recognition deficits observed in patients, NER could therefore be quite relevant for behavioral screening in the context of BD. Finally, we included the NOR test of episodic memory.

In this work, we indeed observed that GBR has an impact on behavioral phenotypes in the SLD and the NER, with a strong switch of responses to threat in the SLD from freezing to active flight in both sexes, and an altered preference for the stressed mouse in the NER in males only. We also replicated the increased anxiety, anhedonia and locomotion obtained in other studies.

## 2. Material and methods

### 2.1. Animals

C57BL/6N male and female mice (12-14 weeks old) were all purchased from Janvier Labs (France). Group composition was as follows: 10 males and 10 females for the GBR treated group, same for the saline injected control group, and 25 males and 25 females that served as emotional targets in the NER. At first, injected mice were socially housed (5 per cage), then were individualized following SLD mice to avoid social fights due to regrouping. Social targets for the REN were kept in groups at all times. All animals were housed under standard housing conditions with a 12/12 light/dark cycle (lights on from 8:30 a.m. to 8:30 p.m.) and food and water ad libitum. All animal care and experimental procedures followed national and European (2010/63/EU) guidelines and were approved by the French Ministry of Research (APAFiS: #49684-2024042511181919_v7).

### 2.2. Drugs

GBR 12909 (16 mg/kg, CliniSciences, France) was prepared in saline solution for a final injection volume of 0.1 ml/10 g body weight, and dissolved after 45 min heating at 60 °C. The animals received daily prepared solutions through intraperitoneal injection 30 minutes before some behavioral tests (Figure 1). Control group received saline injection. Animal were randomly allocated to the experimental groups between cages for the Saline (Sal) vs GBR treatments. To minimize potential confounding factors all cages were located within the same room and rack in the animal facility.

**Figure 1.**
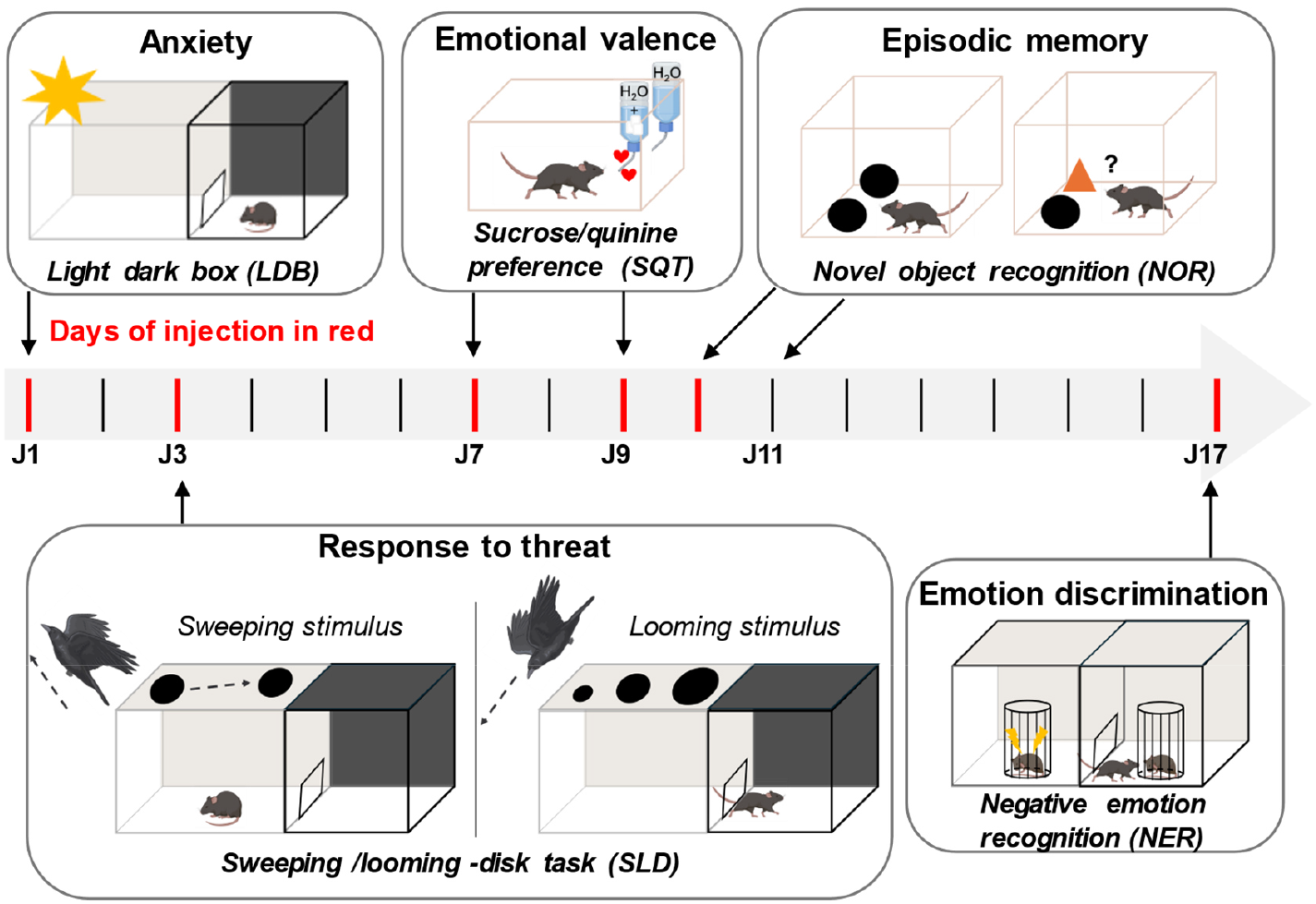
Both the male and the female cohort followed the same experimental schedule over 17 days. Each test evaluates a different symptomatic dimension of BD (top of the boxes). GBR was administered intraperitoneally 30 minutes prior to behavioral tests to characterize the associated mixed manic-like phenotype. Consequently, there were 6 injections in total (on days in red) separated by at least 24 hours, one per behavioral test except for the SQT in which there was one injection right before sucrose presentation, and one right before quinine. Some tests needed prior habituation, hence the intervening days between tests (black).

### 2.3. Behavioral tests

Behavioral tests were performed simultaneously in GBR-treated and saline-injected animals. Due to practical considerations (e.g. space, as well as odor and pheromone communication between males and females), a first cohort of males was performed followed by a cohort of females in the same experimental conditions. Before every behavioral test, mice were habituated ∼30 min to the testing room. To assess GBR effects on males and females, behavioral tests were performed for 18 days (cf Figure 1 for detailed schedule). All the behavioral tests were videorecorded and analyzed by a tracking system (Noldus Ethovision 17, Netherlands). Behavioral arenas were cleaned after each mouse with 70% alcohol to minimize olfactory bias.

#### 2.3.1 Light dark box (LDB)

LDB is an assay commonly used to measure anxiety-like phenotype in mice (Bloch and Belzung, 2023). It is composed by 2 compartments, one dark (20×20 cm) and one brightly lit (∼600 lux, 20×30cm). Mice were first placed in the dark compartment and time spent in compartments, latency to go in light zone and distance were noted for 6 minutes. Lower exploration of the lit compartment is typically interpreted as higher anxiety-like phenotype.

#### 2.3.2. Sweeping looming disk (SLD)

We adapted the SLD based on the original study on differential flight or freezing responses to sweeping or looming disk stimuli (De Franceschi et al., 2016). The test evaluates the reactivity to fear and the avoidance response to a threatening stimulus. The behavioral arena was the same as the light dark box arena. The day before test, mice were habituated to the device during 15 minutes at ∼15 lux, with a computer screen covering the lit compartment displaying the same grey intensity as the one used as background for stimulus presentation. On the next day, after 5 minutes of additional habituation, a total of 6 stimuli was presented separated by at least one minute. Stimulus presentation was triggered by the mouse spending at least one second in the middle of the lit area (live video and launch of stimuli was managed by Noldus Ethovision 17, Netherlands). 3 sweeping stimuli alternating with 3 looming stimuli were presented. The ‘loom’ stimulus is considered to imitate the threatening approach of an aerial predator. It consisted in the sudden appearance of a small black disk (∼1/18^th^ of screen height in diameter) in the center of the screen that stayed on for 3 seconds, then increased to take the full screen (∼60% of screen height in diameter) in 2 seconds and stayed on for three more seconds. The ‘sweep’ stimulus is considered to mimic the lookout of an aerial predator at the prey from a distance. It consisted in a black disk (∼1/14^th^ of screen height in diameter) crossing the screen in a straight diagonal over 4 seconds. Stimuli parameters were inspired by previous work (De Franceschi et al., 2016; Neira et al., 2022) and personal observations.

#### 2.3.3. Sucrose and quinine preference test (SQT)

We followed the methodology used in the recent study on affective bias in the GBR model (Bigot et al., 2022). Mice were habituated to two bottles filled with drinking water overnight. After habituation, mice had the choice between water and a sucrose solution (10g/L) or a quinine solution (0.1 mM) with a 24h break in between with 2 bottles of water. Bottles were weighed at 2h, 6h and 24h. Bottles were swapped after each weight measurement to ensure that the mice did not develop a side preference. Sucrose and quinine preferences were calculated as the percentage (amount of sucrose or quinine solution consumed × 100/total volume consumed) as well as sucrose and quinine intake (volume of quinine or sucrose/animal weight).

#### 2.3.4. Novel object recognition (NOR)

NOR was divided in a learning phase followed by a test phase. The learning phase consisted of placing the mouse into the chamber containing two identical objects for 5 minutes. Following a delay period of 24 hours, mice underwent a 5 min test phase where they were placed in the same chamber containing one of previously encountered objects and a novel object in the same positions as in the learning phase. The test was run in ambient light (∼250 lux) in a 40×40cm arena.

#### 2.3.5. Negative emotion recognition (NER)

NER was performed as previously described by another group (Scheggia et al., 2020). Experimental mice were individually habituated to the arena for 2 consecutive days during 15 minutes at 60 lux. The arena consisted of a standard cage divided into two zones (40×20cm each). In each compartment, a cylindrical cage was placed, preventing direct physical contact but allowing sensory interaction with the social target: olfactory, visual, and auditory. All the demonstrator mice, both “neutral” and “stressed,” were matched to the experimental mice by sex, age, and strain. Neutral mice were naïve, unmanipulated, and kept in their home cages with ad libitum access to food and water. Stressed mice were subjected to gentle restraint for 15 minutes using a restraint tube, immediately before being placed in the cylindrical cage. On the test day, experimental mice were placed in the arena for 6 minutes, and the time spent in each of the two zones as well as in contact with the two targets was measured.

### 2.4. Statistical analysis

Statistical analyses were performed with GraphPad Prism 8 software (USA). Normality was assessed using the Shapiro-Wilk test. Consequently, parametric or non-parametric tests were performed: unpaired Student, one sample t-test, Mann–Whitney, 2 or 3-way repeated measures ANOVA or Mixed-effect model when some values were missing. Tukey’s multiple comparisons test was applied for post-hoc analyses. Log-rank mantle-cox test was used for survival curve analyses, in which significance was corrected for multiple comparisons by the Holm-Bonferroni method (K=8 for figure 5, K=4 for figure S3). As males and females were tested in separate cohorts, cohort and sex effects could not be distinguished. For that reason, we split analyses for males and females. We added analyses including sex as a factor for NER in supplementary data, in which we cannot rule out the cohort effect, although control males and females were generally comparable.

## 3. Results

### 3.1. GBR elicits anxiety, hedonic bias and hyperlocomotion in males

A first goal of our study was to investigate if we would be able to replicate phenotypes associated with the mixed manic-like phase immediately following GBR injection as described previously in males (Bigot et al., 2022), and to characterize those phenotypes in females. In this study it has been shown that within an hour post-injection GBR provokes anxiety-like behaviors, anhedonia, and hyperlocomotion. For that purpose we used a standard anxiety test (Bloch and Belzung, 2023), the light dark box (LDB) that to our knowledge has not been tested in the GBR model to further validate the anxiety-like phenotype observed in other paradigms such as the elevated plus maze and the open field (Bigot et al., 2022). We also replicated the sucrose and quinine preference experiment (SQT) from that study to test for positive and negative emotional biases respectively. In addition, we also used a well-known episodic memory test to identify potential memory alterations in the GBR model.

GBR-treated males had increased anxiety-like phenotypes in the LDB (Fig. 2a,c). Indeed, they both had a higher latency to enter the light compartment and tended to spend less time there, which is commonly interpreted as a higher-level of anxiety-like behaviors. In contrast, females-treated did not show differences from saline-injected controls neither in latency to enter nor in time spent in the light zone (Fig. 2b,d). Likewise, GBR-treated males exhibited an anhedonic-like response in the SQT, with an overall decreased sucrose preference and intake compared to controls (Fig. 3a,c). In females, GBR had no significant effect on sucrose preference or intake (Fig. 3 b,d). In addition, there was no effect of GBR on quinine preference or intake in both sexes, suggesting no change in the drive for aversive stimuli was associated with GBR treatment (Fig. 3e-f).

**Figure 2.**
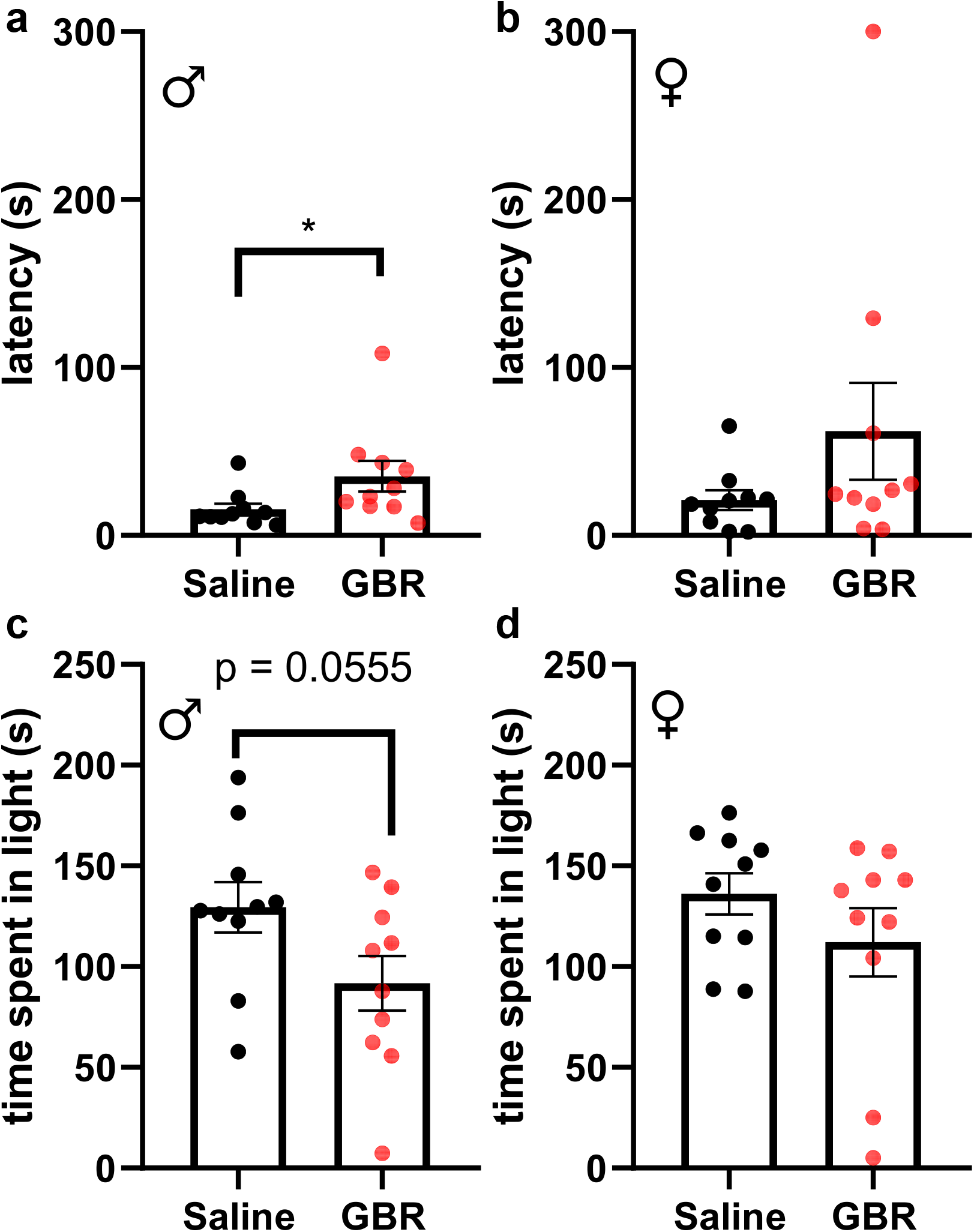
LDB shows anxiety-like phenotypes in males but not in females. (**a**) Males showed a reduced latency to enter in the lit compartment (Mann Whitney U=18, *p*=0.0147) (**b**) but not females (Mann Whitney U=31.50, *p*=0.1717). (**c**) Likewise, time spent in the light compartment was nearly significantly higher in males (unpaired t-test, t=2.047, df=18, *p*=0.0555) (**d**) while not in females (Mann Whitney U=37, *p*=0.3527). Significance key: \**p*<0.05.

**Figure 3.**
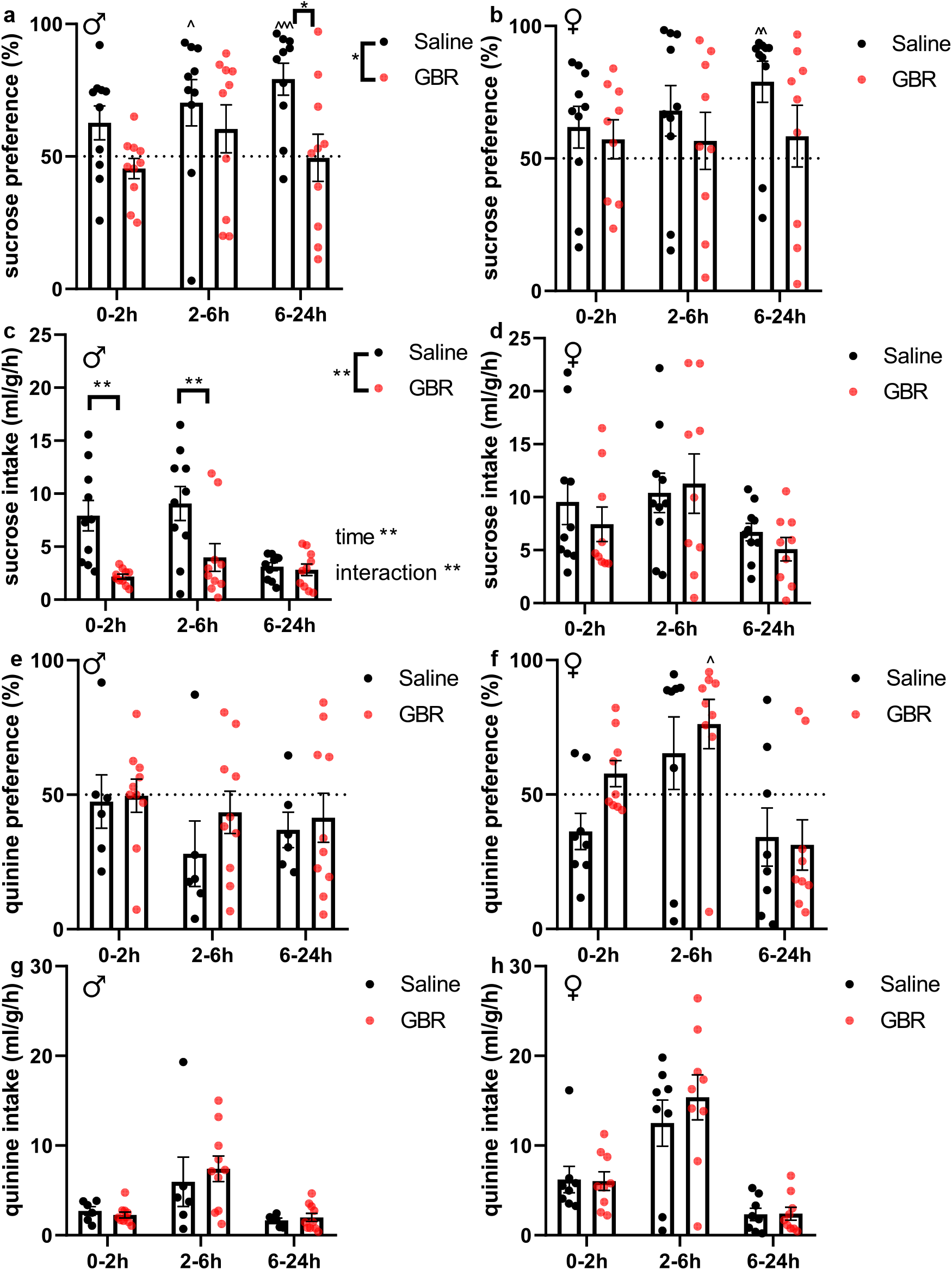
Unlike females, GBR males show anhedonia-like reduction in sucrose preference. (**a**) Sucrose preference was reduced in males (2-way repeated measures ANOVA, treatment F_(1,18)_ = 7.819, *p*=0.0119, time F_(1.553,27.95)_ = 1.624, *p*=0.2170, interaction F_(2,36)_ = 1.047, *p*=0.3613, followed by Sidak’s multiple comparison test). (**b**) Sucrose intake was also diminished in GBR males (2-way repeated measures ANOVA, treatment F_(1,18)_ = 11.91, *p*=0.0028, time F_(2,36)_ = 12.35, *p*=0.0017, interaction F_(2,36)_ = 5.296, *p*=0.0096, followed by Sidak’s multiple comparison test, *p*=0.0011 and *p*=0.0041 at 0-2h and 2-6h respectively. (**c**) There was no significant difference between groups in females for preference (2-way repeated measures ANOVA, treatment F_(1,17)_ = 1.191, *p*=0.2903, time F_(1.633,27.96)_ = 1.272, *p*=0.2902, interaction F_(2,34)_ = 0.9281, *p*=0.4051) (**d**) or intake (2-way repeated measures ANOVA, treatment F_(1,17)_ = 0.2724, *p*=0.6085, time F_(1.389,23.61)_ = 4.782, *p*=0.0284, interaction F_(2,34)_ = 0.4984, *p*=0.6119). GBR had no effect neither on quinine preference in males (**e**) or females (**f**), nor on quinine intake (males in (**g**), females (**h**)). As for sucrose preference, 2-way repeated measures ANOVA followed were performed and detailed statistics are available upon request. Significance key: between group comparison, \**p*<0.05, ** *p*<0.01; one group within a timepoint comparison to chance, ^*p*<0.05, ^^*p*<0.01, ^^^*p* <0.001 (one sample t-tests, detailed statistics available upon request).

Regarding episodic memory, we could not demonstrate the presence of a novel object memory in control animals. Neither males nor females (Fig. S1a,b) showed a difference between the familiar and the novel object on day 2 (see discussion for possible methodological issues). Nevertheless, GBR administration decreased object exploration overall in both sexes (Fig. S1a,b) and increased locomotion in males and in females (Fig. S1c,d).

In conclusion, we replicated the heightened anxiety, hyperlocomotion and decreased hedonia (in our hands only towards the innately positive valence stimulus, sucrose) previously observed in the GBR model in male animals. Intriguingly, females did not show any significant anxiety-like or anhedonia-like phenotypes.

### 3.2. GBR modifies behavioral responses to threatening stimuli in both sexes

Building on reports in the last decade on adaptative and ecological fear responses to threatening stimuli presented on a screen in mice (De Franceschi et al., 2016; Yilmaz and Meister, 2013) as well as on the recent interest for these behaviors in other fields of neuropsychiatry (Neira et al., 2022), we adapted a sweeping looming disk (SLD) procedure consisting of 3 presentations of a sweeping disk alternating with 3 presentations of a looming disk (see methods for details). The interest of having two types of stimuli is that they elicit two types of responses: an active flight response typically for a fast imminent threat such as the looming disk, or a freezing response for a potential threat such as the sweeping disk. The proportion of freeze and flight has been shown to be influenced by the speed of the stimulus and the characteristic of the area (De Franceschi et al., 2016). We opted for a moderate speed for both stimuli because in our hands faster stimuli did not elicit freeze or flight as efficiently (personal observation). Thus, we were interested in the effect of GBR on fleeing and freezing according to stimulus type in both sexes.

When observing only the first behavioral response, male controls do not privilege freezing or fleeing in response to either stimulus (Fig. 4a,c). GBR treatment increases flight in response to both stimuli and promotes flight over freezing in response to both stimuli. Female controls have an overall different profile (Fig. 4b,d): they flee very little to both stimuli, but strongly prefer to freeze in response to sweep (above 85% of freezing responses to sweep after 4 seconds). GBR injection in female yields globally similar responses as in males, with a notably strong decrease in freezing responses to sweeping stimuli and overall increase in flight to both stimuli. To better apprehend this switch of responses due to GBR, we represented the first response to each threat in the chronological order of trials (Fig. 4e). Saline animals mainly responded by freezing to threatening stimuli, while GBR induced a complete change of strategy in both sexes, GBR-treated animals strongly preferred flight as a first response. We also compared all responses, regardless of first occurrence (Fig. S3). In other words, we included both freezing after a flight, and flight after a freezing epoch (‘freeze and flight’). This complementary analysis yielded similar results, GBR increasing flight and decreasing freezing in both sexes.

**Figure 4.**
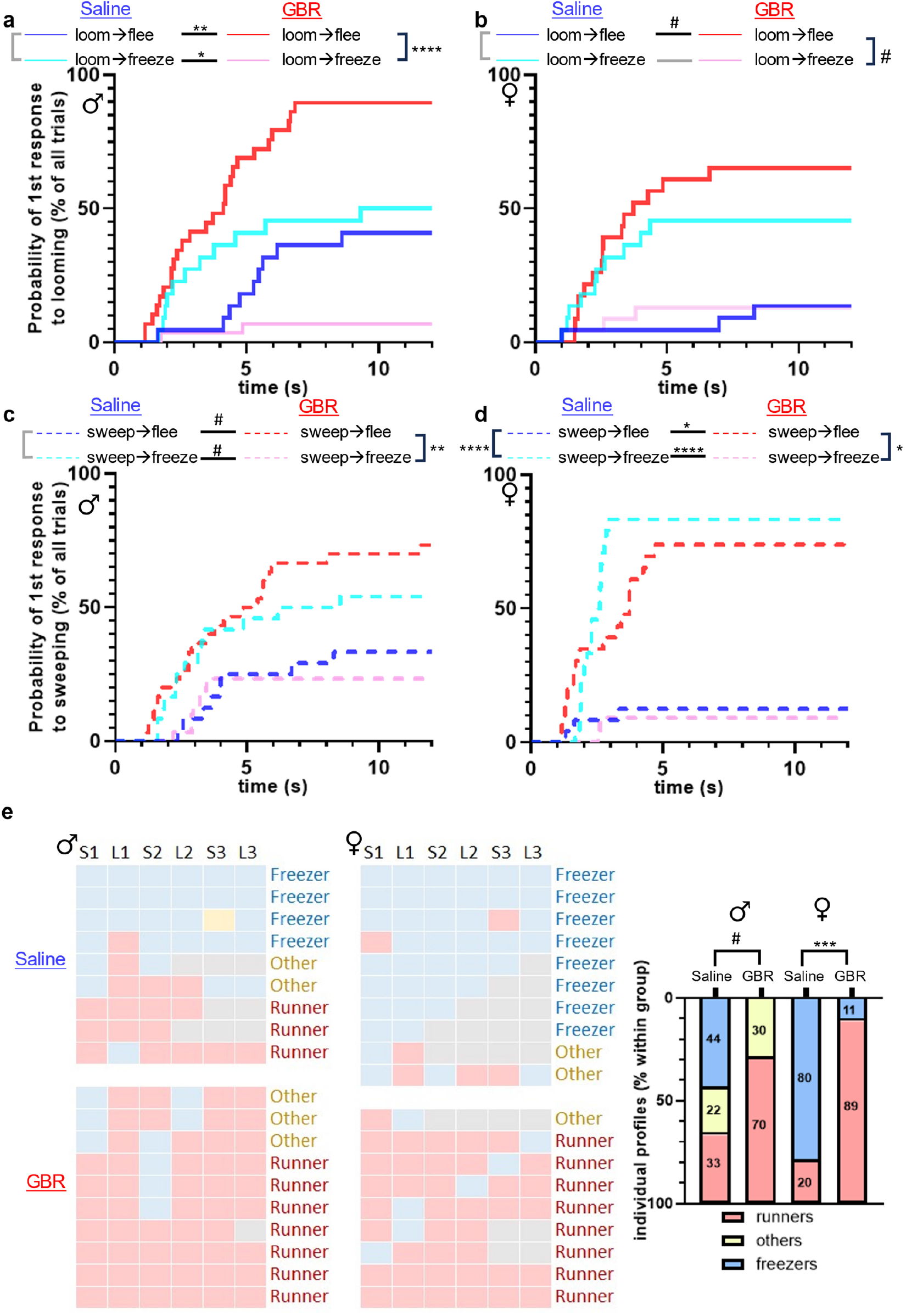
These graphs show the preferred response to each threat (**a**,**b** for looming, **c**,**d** for sweeping) by plotting only the first response (freeze or flee) for males (**a**,**c**) and females (**b**,**d**). Below the graphs are shown legends as well as main effects as black arrows pointing in the direction of the higher latency. (**a**) GBR increases fleeing and decreases freezing in males (for GBR/saline flight in response to loom, χ2=13.54, *p*=0.0014, GBR/saline freezing in response to loom, χ2=9.119, *p*=0.0125, within GBR flight/freeze comparison, χ2=37.92, *p*<0.0001). (**b**) Females showed similar trends, GBR tending to increase flight in response to looming (for GBR/saline loom flight, χ2=6.140, *p*=0.0528, for withing GBR flee/freeze comparison to loom, χ2=5.434, p=0.0591). (**c**) As for looming stimuli, GBR tended to increase flight and reduce freezing in males (for GBR/saline flight in response to sweep, χ2=4.977, *p*=0.0771, GBR/saline freezing in response to sweep, χ2=5.768, *p*=0.0652, within GBR flight/freeze comparison, χ2=11.87, *p*=0.0036). (**d**) Interestingly, female saline controls had a strong preference for freezing over fleeing (χ2=11.87, *p*<0.0001). GBR induced similar changes as in males by strongly reducing freezing (χ2=21.89, *p*<0.0001) and increasing flight (compared to control χ2=8.718, *p*=0.016, within GBR group χ2=9.707, *p*=0.0108). Survival analyses were performed using log-rank Mantel-Cox tests and comparing the curves two by two with *p*-values reaching significance adjusted for multiple comparisons (K = 8). Trends displayed all passed significance uncorrected, only corrected *p*-values are shown. (**e**) The previous observations were confirmed and completed by analyzing the first response for each individual mouse at each trial. Blue squares denote freezing first, red fleeing first. Some individuals did not complete all trials, grey squares denote trials not triggered. A threshold of 12 seconds was applied, meaning that if latency exceeded 12 seconds for both response type, the trial was considered a time out (yellow square, only one occurrence). One can see a major effect of GBR in both groups, in which it promotes flight as a first response instead of freezing. Individuals were classified according to their response profiles: ‘freezers’ and ‘runners’ responded with freezing and flight respectively in most trials (at least 75% of one response type). The rest of the animals without a clearly preferred response were classified as ‘others’. Contingency analysis demonstrates a significant effect of GBR on response profile in both sexes (χ2=5.763, *p*=0.0560 in males, χ2=15.43, *p*=.0004 in females). Significance key: ^#^*p*<0.1, \**p*<0.05, \*\**p*<0.01, \*\*\**p*<0.001, \*\*\*\**p*<0.0001.

We also wondered if the sequential presentation of threatening stimuli could induce habituation, but there was no significant effect of stimulus repetition (Fig. S2). In both males and females, GBR importantly increases flight response for both sweeping and looming stimuli (Fig. S3a,b). GBR-treated males have a decreased freezing response to looming stimulus and a trend in the same direction for sweeping (Fig. S3c). GBR further decreases both freezing and looming to both stimuli in females (Fig. S3d).

In conclusion, GBR increases active responses to a threat (flight) while it decreases passive responses (freezing). More specifically, males and females show differences in baseline responses to threat, notably with females strongly preferring freezing to sweeping stimuli. GBR abolishes those differences by switching the response strategy to active flight responses to any threatening stimuli.

### 3.3. GBR alters emotional discrimination in a sex dependent manner

A relevant phenotypic dimension pertaining to emotion regulation and empathic-like behaviors is emotion recognition. A recent study (Scheggia et al., 2020) has developed an emotion discrimination paradigm in mice based on the tendency to favor interaction with a stressed conspecific. The authors used it to investigate the neurobiological underpinnings of this behavior, with no clear baseline difference between sexes. Given its relevance for BD, we used the same behavioral paradigm focusing on preference to interact with a control mouse or a stressed mouse (negative emotion recognition, NER). As for the LDB, GBR elicited hyperlocomotion in males (Fig. S4a) and there was a similar trend in females (Fig. S4b). Both in males and females, saline injected controls spent more time in contact with the stressed target (Fig. 5a,b). In addition, saline males spent more time in the stressed target compartment (Fig. 5c), which was not observed in saline females (Fig. 5d). Interestingly, GBR affected sexes differentially regarding emotion discrimination. In GBR males, it decreases both time spent in contact and in the compartment of the stressed target compared to controls, and GBR males lose the preference for the stressed target compared to chance (Fig. 5a,c). In contrast, GBR females do not differ from saline females significantly regarding contacts and time spent in the stressed target zone and spend both more time in contact and in the compartment of the stressed target (Fig. 5b,d).

**Figure 5.**
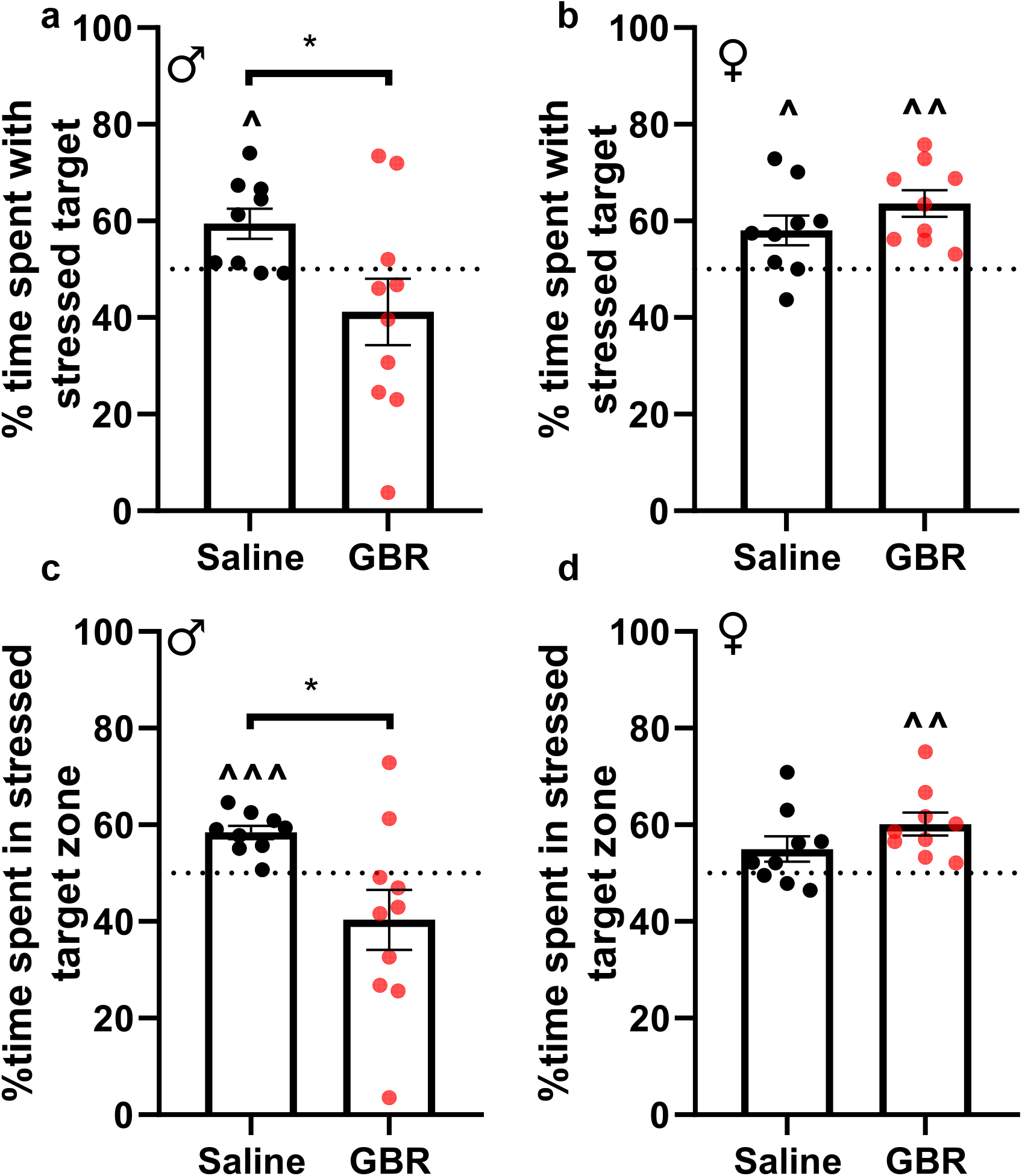
GBR males and females exhibit different performances in emotion discrimination. (**a**) GBR males spent less time in nose contact with the emotional target than saline (unpaired t-test t=2.323, df=17, *p*=0.0328). Saline males spent more time to interact with the stressed target than chance (50% of interaction with each target), unlike GBR-treated animals (one sample t tests, t=3.0131, df=8, p=0.0163, t=1.280, df=9, p=0.2324 respectively). (**b**) In contrast both GBR (t=4.97 df=8, p=0.0011) and saline (t=2.605, df=8, p=0.0314) females contacted more the emotional target than the control target and there was no significant difference between the two groups (unpaired t-test t=1.354, df=16, *p*=0.1947). (**c**) Time spent in each zone yielded similar results, with saline spending more time in the emotional target compartment while GBR showing no significant preference (one sample t tests, t=6,013, df=8, p=0.0003, t=1,562, df=9, p=0.1526 respectively, unpaired t-test, t=2.711, df=17, *p*=0.0148). (**d**) As for females, the GBR group preferred the compartment where the stressed target was located (t=4.263 df=8, p=0.0028) while controls showed a similar but not significant trend (t=1.907 df=8, p=0.093). However, the two groups did not differ significantly (t=1.462, df=16, *p*=0.1632). Significance key: between group comparison, \**p*<0.05; one group comparison to chance, ^*p*<0.05 ^^*p*<0.01.

To explore this sex effect further (in spite of the confounding cohort factor, cf methods), we ran a 2-way mixed ANOVA including sex for nose contact and time in zone (Fig. S4). For the former, there was an interaction between treatment and sex (Fig. S4c). In males there was a reduction in nose contact with the emotional target, which was not observed in females (as in Fig. 5a,b). For the time spent in each zone, there was a treatment and sex interaction as with the same group differences as for nose contact (Fig. S4d).

In brief, saline control mice showed interest in the stressed mouse comparable in both sexes. GBR decreased social interaction with a stressed conspecific in males, while it did not have a visible effect in females.

## 4. Discussion

In this work, we investigated different behavioral dimensions in male and female animals during the mixed manic-like phase following GBR injections. We introduced two novel assays (SLD and NER) that had never been used in preclinical models of BD. Importantly, we found that there were highly sensitive to the GBR-induced manic-like state in both male and female mice. Moreover, behavioral reactions to GBR differed according to sex for certain measures. As hypothesized, GBR-treated animals showed a change of behavioral strategy in response to threat in both sexes, while preference for a stressed conspecific was abolished only in males. This study therefore validates SLD and NER as novel behavioral tools to assess BD-related symptomatic dimensions that are impacted in patients. Furthermore, this adds validity to the GBR mania model, since altered risk-assessment, hypervigilance, and negative emotion recognition are observed in manic patients.

In the SLD, GBR globally increased flight and decreased freezing in both sexes. However, looking at the first prioritized between flight and freeze revealed a sex difference in the baseline strategy to respond to threat, saline females showing a strong preference for freezing to sweeping stimuli while males did not favor flight or freeze whatever for either stimulus. For the latter, this is in agreement with published results for a looming stimulus of moderate speed in C57BL6 male mice (De Franceschi et al., 2016). GBR changes these priority allocations by globally promoting flight over freezing regardless of stimulus type. In other words, GBR has a major impact on responses to both imminent and distant visual threats by increasing flight and decreasing freezing responses (Table 1). In the NER, GBR provoked a reduction in interaction with the stressed animal in males but not in females. Further experiments including more animals would be needed to determine if GBR accentuates this preference or merely does not affect this phenotype in females.

**Table 1.** Summary of main findings. GBR administration affects both sexes, but in a different fashion. Anxiodepressive phenotypes as measures by the LDB and anhedonia are affected by GBR in males, but less (or none) in females. The SLD demonstrates an overall increase in flight responses and an associated decrease in freezing to both types of stimuli. More particularly, males and females show a different strategy regarding first response, GBR-treated males favoring flight to looming stimuli while females decrease freezing to sweeping stimuli and have an increased flight to this stimulus type. In the REN, GBR-treated males show an opposite phenotype to controls, with less interest for the emotional mouse, while GBR-treated females keep a pronounced interest in the stressed target.

| Compared to saline | Anxiety | Anhedonia | Locomotion | Reaction to threat | Emotional preference |
| --- | --- | --- | --- | --- | --- |
| ♂ GBR | ↑ | ↑ | ↑ | ↑ flight<br>↓ freezing | ↓ |
| ♀ GBR | ns | ns | ↑ | ↑ flight<br>↓ freezing | Not affected |

We also evaluated anxiety-like phenotypes and hedonic bias in the LDB and SQT respectively to replicate previous result in the GBR mania model (Bigot et al., 2022; Young et al., 2010) and to explore its effects in female mice. Our results confirm an increase in anxiety-like phenotypes, locomotion and decreased sucrose consumption (anhedonia-like) in males, as previously reported (Bigot et al., 2022), but not in females. However, we could not replicate the increase in quinine preference. This discrepancy in anxio-depressive phenotype between male and females has been reported elsewhere (Kokras and Dalla, 2014).

We also attempted to study potential episodic memory deficits caused by GBR during memory encoding (object learning) using a the widely used NOR paradigm. Neither saline nor GBR treated animals showed any preference for the novel object on the second day, which prevented the study of any potential memory alteration due to GBR. There are several variations of this task that could permit to improve conditions for learning (Leger et al., 2013), for instance by introducing an additional day of habituation and performing the task under dim light. This could help mitigate the stress due to the novel environment and allow learning in the object presentation phase.

These data support the value of characterizing emotion dysregulation for the study of BD mood states. In addition to anxiety-related behaviors and anhedonia in the SQT, we believe NER to be a valuable addition to assess emotion perception in conspecifics in models of psychiatric disorders. In this context, SLD also appears to be a robust method to quantify innate reactions to a threat. These two dimensions are especially relevant to BD symptomatology, as patients show alterations in empathy and emotion recognition (Russo et al., 2015), as well as maladaptive responses to stress (Muhtadie and Johnson, 2015). They also show altered startle responses (Mao et al., 2019).

The behavioral alterations observed here are also in line with the powerful dopamine selective reuptake inhibition of GBR that affects dopamine transmission widely (Andersen, 1989; Irifune et al., 1995). This translates into hyperlocomotion (Queiroz et al., 2015), anhedonia, anxiety, (Bigot et al., 2022), impulsivity (van Enkhuizen et al., 2013) and increased attention (Loos et al., 2010). These behavioral dimensions are respectively modulated by dopamine release in various brain regions including striatum (including nucleus accumbens), amygdala and medial prefrontal cortex (Costa and Schoenbaum, 2022; Stanton et al., 2019). Our study adds response selection to threat to these dimensions with the SLD, also known to be regulated by dopaminergic release to the superior colliculus (Bolton et al., 2015; Montardy et al., 2022).

Interestingly, innate responses to fast visual and auditory stimuli interpreted as threats are respectively processed in the mammalian tectum in the superior (Isa et al., 2021) and inferior (Nobre et al., 2003) colliculi, which project to the amygdala forming the ‘short’ path to this major integrator of fear and emotion-related cognition. In humans, a similar neuroanatomical organization of these short routes to amygdala has also recently been demonstrated, and likewise are involved in fast response to threats in both sensory modalities (Rafal and Koller, 2025). This tectal pathway has also been implicated in emotion recognition in humans (Méndez et al., 2022), and has been involved in other neurodevelopmental and psychiatric disorders such as autism (Huang et al., 2022), schizophrenia (Kang et al., 2008) and post-traumatic stress disorder (Olivé et al., 2018). In the future, the dissection of the interplay between this ‘short’ pathway and thalamo-cortical emotion processing in response to the SLD and NER could be a novel avenue to study the neurobiology of BD states in preclinical models.

## Legends

**Figure S1.**
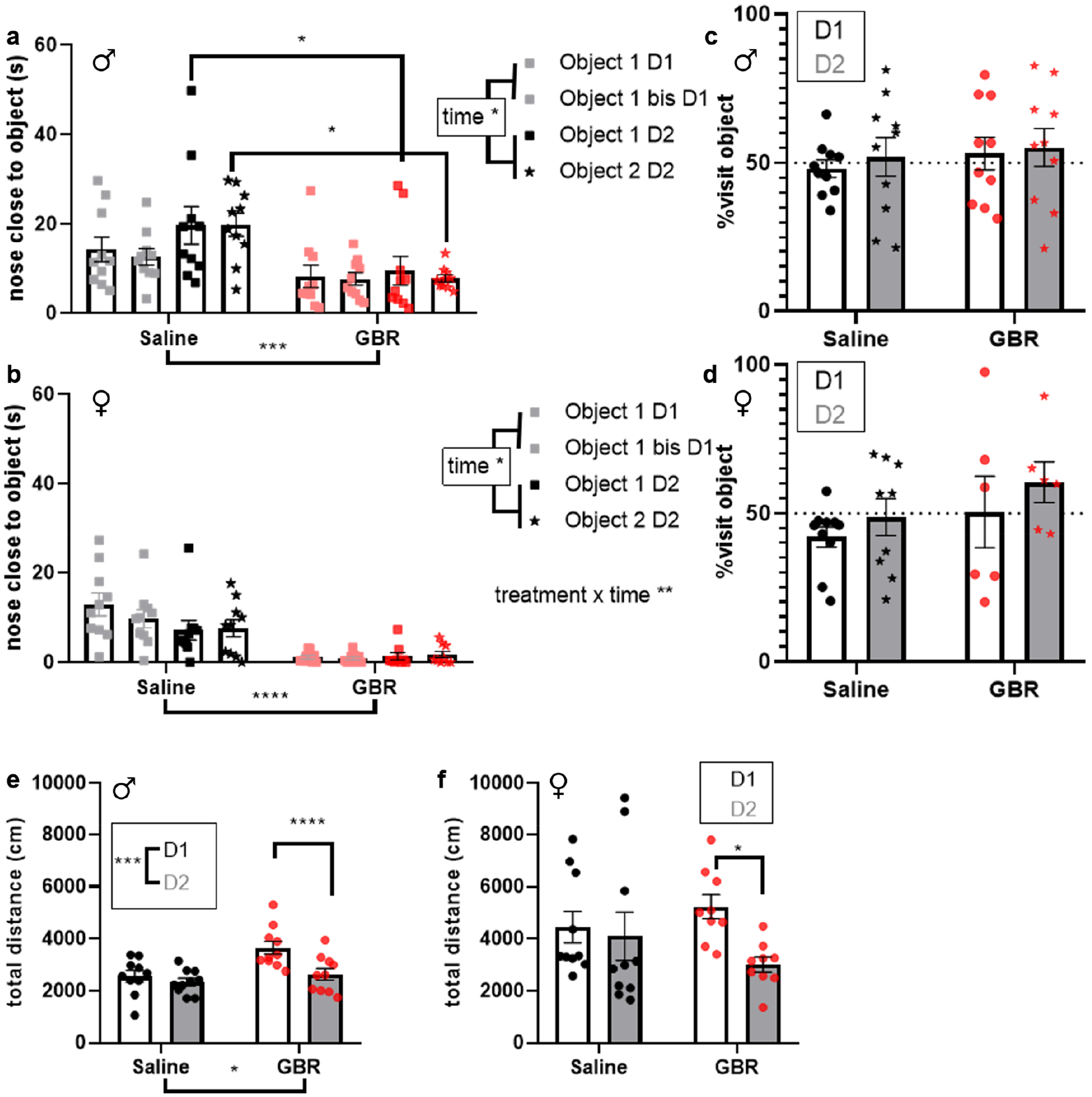
GBR-treated animals have a decreased interest for all objects, albeit no difference in learning can be shown in the NOR. (**a**) Male and (**b**) female GBR-treated mice globally spend less time in contact with the objects, regardless of object novelty introduced in day 2 (object 2 in red on the graphs, for males, 3-way ANOVA, treatment F_(1,36)_ = 14.88, *p*=0.0005, time F_(1,36)_ = 5.720, *p*=0.0221, novelty F_(1,36)_ = 0.1899, *p*=0.6656, followed by Sidak’s multiple comparison test, for females, 3-way ANOVA, treatment F_(1,34)_ = 27.60, *p*<0.0001, time F_(1,34)_ = 6.564, *p*=0.015, novelty F_(1,34)_ = 0.1823, *p*=0.6721). (**c**,**d**) Indeed, there was no difference in preference compared to chance or between groups for either same objects on day 1 or a novel object (star) on day 2 (2-way ANOVAs for both sexes, detailed statistics available upon request). (**e**) Additionally, GBR induced hyperlocomotion in males on the day of injection (2-way repeated measures ANOVA, treatment F_(1,18)_ = 6.545, *p*=0.0197, time F_(1,18)_ = 22.52, *p*=0.0002, interaction F_(1,18)_ = 9.335, *p*=0.0068, followed by Sidak’s multiple comparison test). (**f**) This effect on locomotion followed the same pattern in females although they was only a trend for a time effect mainly due to the GBR group (2-way repeated measures ANOVA, treatment F_(1,34)_ = 0.06018, *p*=0.8077, time F_(1,34)_ = 4.078, *p*=0.0514, interaction F_(1,34)_ = 0.2.155, *p*=0.1513, followed by Sidak’s multiple comparison test *p*=0.0432). Significance key: \**p*<0.05, \*\**p*<0.01, \*\*\**p*<0.001, \*\*\*\**p*<0.0001.

**Figure S2.**
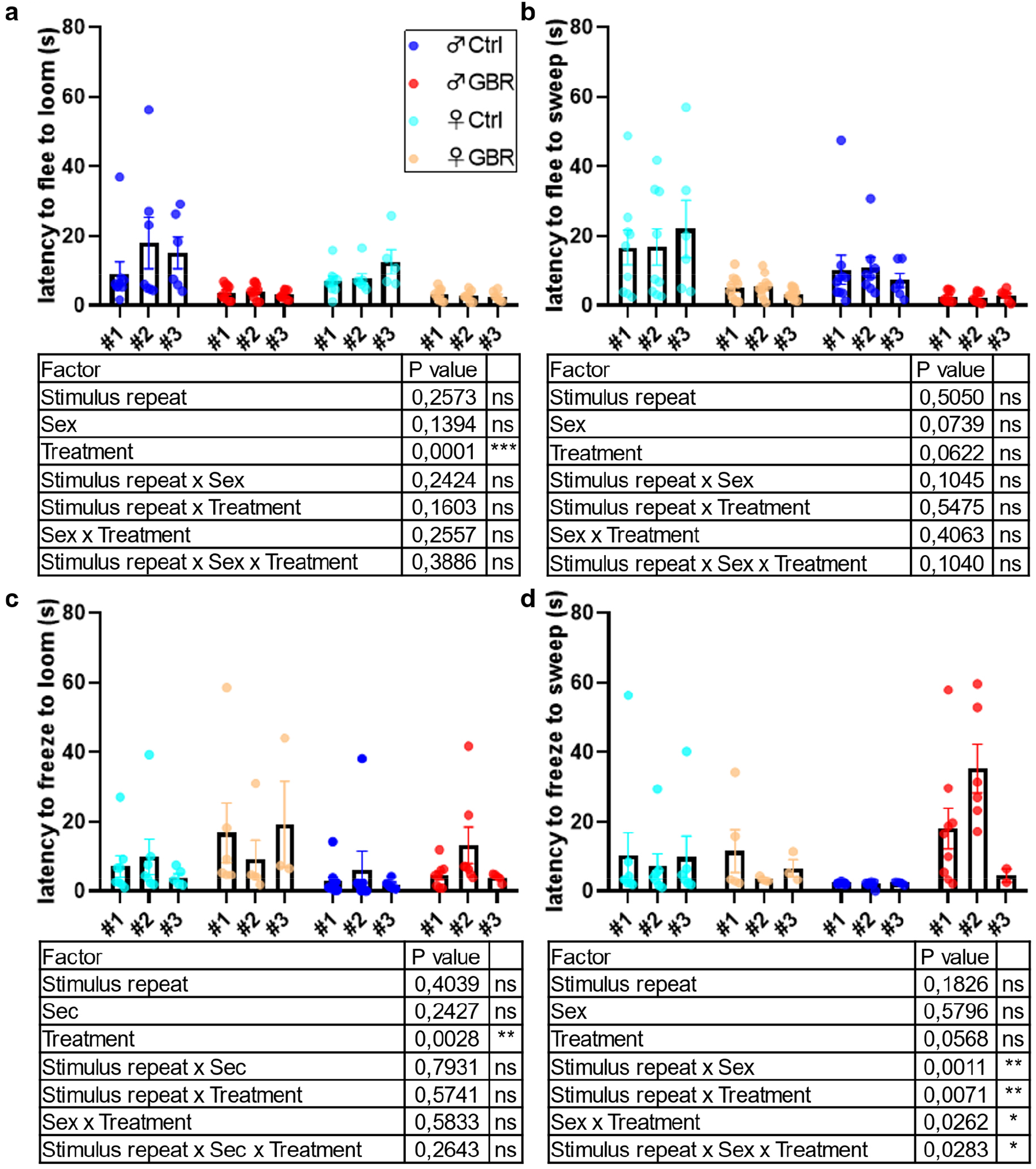
Simulus repetition had no effect on latency to freeze (**a**,**b**) or flee (**c**,**d**) in response to looming (**a**,**c**) or sweeping stimuli (**b**,**d**). For the four situations, 3-way mixed effects analyses were performed with stimulus repetition as a matching factor. Main effects and interactions are summarized in each table below. Detailed statistics are available upon request. Significance key: \**p*<0.05, \*\**p*<0.01, \*\*\**p*<0.001.

**Figure S3.**
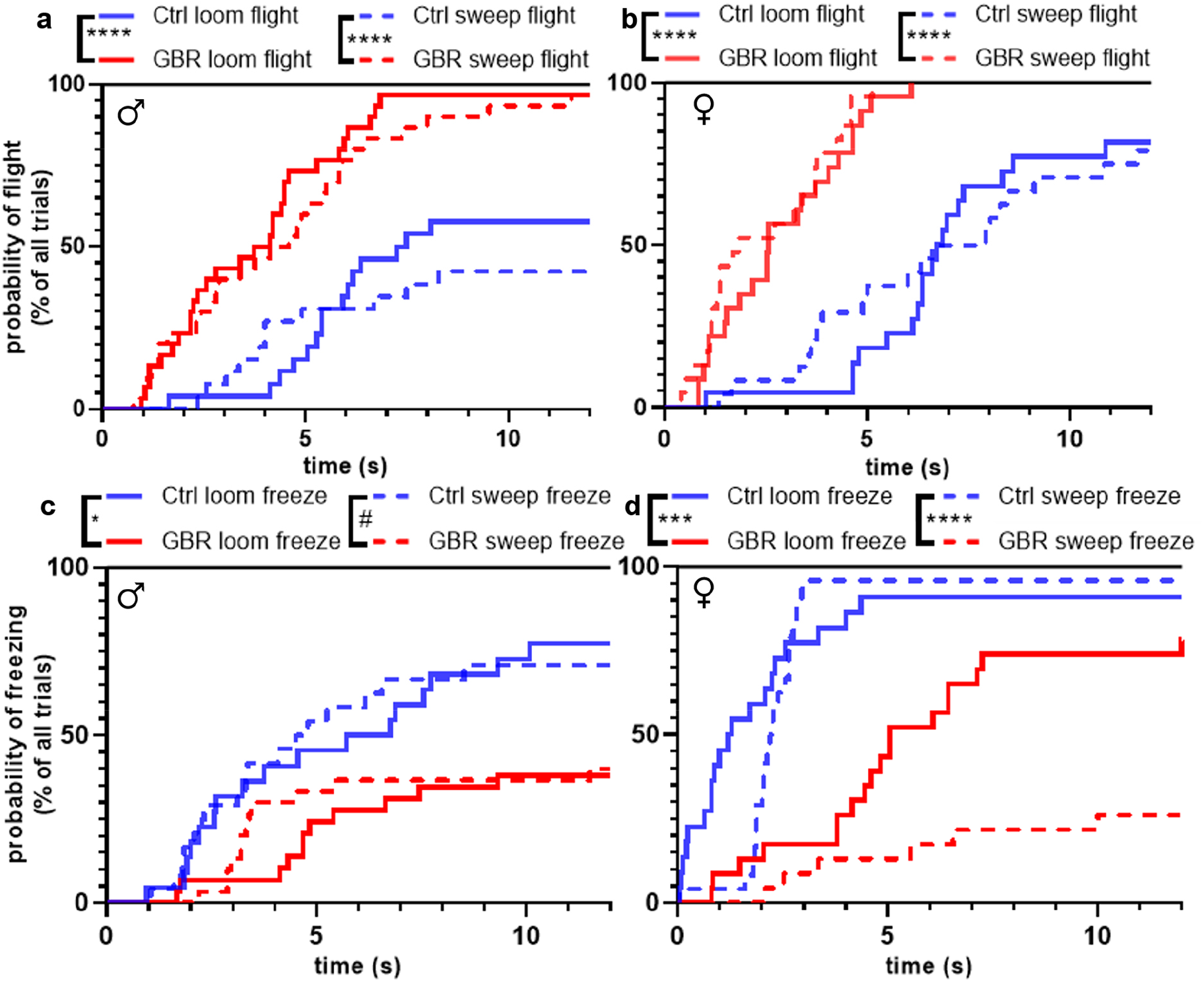
Both males and females treated with GBR globally increase flight and decrease freezing in response to both stimuli. (**a**) GBR-treated males fled more in response to looming stimuli (full line) and sweeping stimuli (dotted line). Survival analysis was performed using log-rank Mantel-Cox tests and comparing the curves two by two with *p*-values reaching significance adjusted for multiple comparisons (K = 4). For GBR/saline flight in response to loom, χ2 = 21.12, *p*<0.0001, for GBR/saline flight in response to sweep χ2 = 22.99, *p*<0.0001, for within GBR group comparison between sweep and loom for flight χ2 = 0.6060, *p*=0.4363, for within saline group comparison between sweep and loom for flight χ2 = 0.06997, *p*=0.7914). (**b**) Similar heightened flight responses were observed in GBR females (for GBR/saline flight in response to loom, χ2 = 37.35, *p*<0.0001, for GBR/saline flight in response to sweep χ2 = 27.66, *p*<0.0001, for within GBR group comparison between sweep and loom for flight χ2 = 0.8347, *p*=0.3609, for within saline group comparison between sweep and loom for flight χ2 = 0.04809, *p*=0.8264). (**c**) GBR-treated males decreased freezing to looming stimuli and also tended to for sweeping (for GBR/saline freezing to loom, χ2 = 8.793, *p*=0.012, for GBR/saline freezing to sweep χ2 = 5.671, *p*=0.0519, for within GBR group comparison between sweep and loom for flight χ2 = 0.02802, *p*=0.8671, for within saline group comparison between sweep and loom for flight χ2 = 0.03207, *p*=0.8579). (**d**) Female controls showed high level of freezing to both stimuli, which was importantly decreased when GBR was administered (for GBR/saline freezing to loom, χ2 = 14.70, *p*=0.0008, for GBR/saline freezing to sweep χ2 = 27.61, *p*<0.0001, for within GBR group comparison between sweep and loom for flight χ2 = 5.955, *p*=0.0294, for within saline group comparison between sweep and loom for flight χ2 = 0.5812, *p*=0.4458). Significance key: ^#^p=0.0519, \**p*<0.05, \*\*\**p*<0.001, \*\*\*\**p*<0.0001.

**Figure S4.**
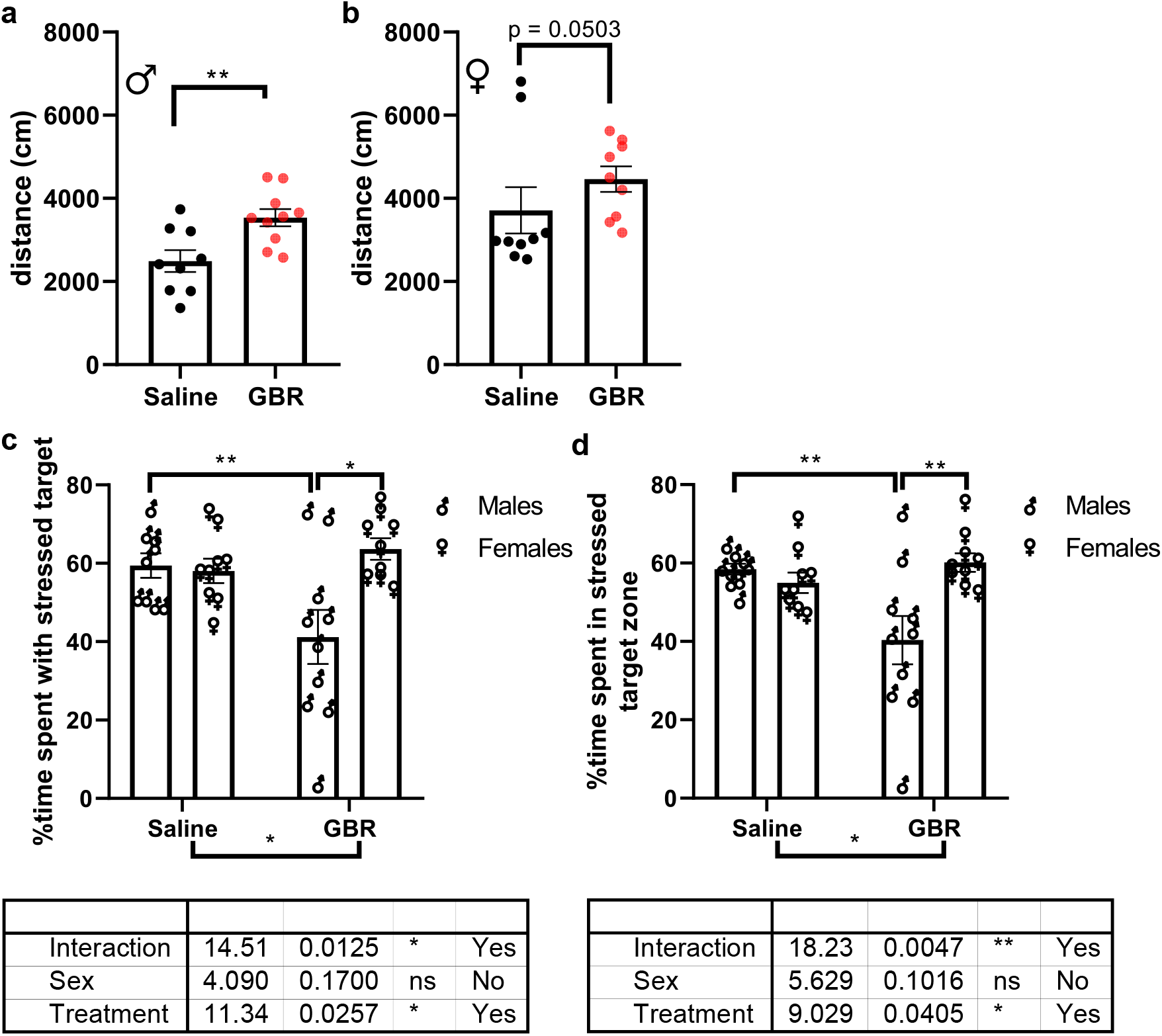
Additional analyses regarding the NER. (**a**) males ran more distance in the arena while females (**b**) showed a similar trend. 3-way ANOVA were performed for comparing directly the two cohorts (males and females) for the time spent with the nose in contact with targets (**c**) and time in each compartment (**d**). They were followed by Sidak’s multiple comparison test for subgroup comparison. Main effects and interactions are summarized in each table below. Detailed statistics are available upon request. Significance key: \**p*<0.05, ** *p*<0.01, \*\*\**p*<0.001, \*\*\*\**p*<0.0001.

